# Persistence of traditional and emergence of new structural drivers and factors for the HIV epidemic in rural Uganda; a qualitative study

**DOI:** 10.1101/516054

**Authors:** Francis Bajunirwe, Denis Akakimpa, Flora P. Tumwebaze, George Abongomera, Peter N. Mugyenyi, Cissy M. Kityo

## Abstract

**Background:** In Uganda, the HIV epidemic is now mature and generalized. Recently, there have been reports of resurgence in the incidence of HIV after several years of successful control. The causes for this resurgence are not clear but suspected to be driven by structural factors that influence large groups of people rather than individuals. The aim of this study was to describe the structural drivers of the HIV epidemic and inform the next generation of interventions.

**Methodology:** We conducted a total of 35 focus group discussions in 11 districts in Uganda. Focus groups consisted of men and women including opinion leaders, civil servants including teachers, police officers, religious, political leaders, shop keepers, local residents and other ordinary persons from all walks of life. The qualitative data were transcribed and analyzed manually. Texts were coded using a coding scheme which was prepared ahead of time but emerging themes and codes were also allowed.

**Results:** Our data indicated there is persistence of several structural drivers and factors for HIV in rural Uganda. The structural drivers of HIV were divided into three categories: Gender issues, socio-cultural, and economic drivers. The specific drivers included several gender issues, stigma surrounding illness, traditional medical practices, urbanization, alcohol and substance abuse and poverty. New drivers arising from urbanization, easy access to mobile phone, internet and technological advancement have emerged. These drivers are intertwined within an existing culture, lifestyle and the mixture is influenced by modernization.

**Conclusion:** The traditional structural drivers of HIV have persisted since the emergence of the HIV epidemic in Uganda and new ones have emerged. All these drivers may require combined structural interventions that are culturally and locally adapted in order to tackle the resurgence in incidence of HIV in Uganda.

## Background

The HIV epidemic in Uganda and many countries in sub Saharan Africa is now mature and generalized. In the late 1980’s and early 1990’s, Uganda experienced a sharp increase in the incidence of HIV cases, but due to several concerted prevention efforts, followed by behavior change, HIV incidence reduced significantly [1–3]. Uganda was the first country to register success in reducing number of new cases of HIV. However, the fortunes reversed and studies started to show that HIV incidence was no longer falling [4]. Studies show the incidence is in fact very high in some groups such as fishing populations [5, 6], which spills over into the general population [7]. The factors responsible for this negative trend are poorly understood.

The potential reasons for this resurgence in HIV incidence may due to structural factors and drivers, which are typically not amenable to individual level interventions [8]. Structural factors and drivers for HIV may be collectively defined as elements, other than knowledge or awareness that influence risk and vulnerability to HIV infection, though drivers specifically refer to a situation where an empirical association within a target group has been established [9]. There is evidence that levels of knowledge on HIV are generally high among countries with high HIV burden, but despite this, structural factors, which are complex and entrenched in the moral, social and cultural fabric of society strongly influence sexual behavior [10–12], often disregarding associated risk of HIV acquisition. These structural drivers and factors have also been broadly defined by some authors as “core social processes and arrangements, reflective of social and cultural norms, values, networks, structures and institutions” [13]. Uganda has had several years of HIV prevention programs, however anecdotal evidence suggests structural drivers and factors for HIV transmission have persisted.

Widow cleansing and inheritance [14–18], and stigma [19, 20] have been well documented as drivers for HIV transmission While these and other drivers are well known, it is not clear if they have persisted and may therefore be potentially responsible for the rise in incidence of HIV in Uganda. In this regard, they have not been well characterized in this resurgence of the epidemic. More studies need to be done to identify and characterize these factors and drivers, and inform the design HIV prevention approaches that target them. Typically, these factors are difficult to tackle, require wider scale community engagement. Interventions at individual level in this instance may not be beneficial, and instead more complex approaches to delivering interventions are required. Therefore, a thorough understanding of these factors is important.

To design interventions targeting these structural factors, researchers and policy makers also need to understand the challenges faced in addressing these drivers. The information will be used to design culturally sensitive interventions. Therefore, the aim of this study was to determine the structural drivers of HIV transmission in rural Uganda and the potential challenges in addressing these drivers.

## Methodology

### Data collection

We conducted a total of 35 focus group discussions (FGDs) in 11 districts in northern, eastern and central Uganda. We designed an FGD guide to identify structural drivers of the HIV epidemic, but also to explore the extent of the already known factors. The FGDs comprised of 8-12 community members and leaders, who were considered to have some knowledge on societal issues related to HIV prevention and transmission. The FGDs were organized along the lines of members that comprised men and women including local council leaders, opinion leaders, civil servants such as teachers and police officers, shop keepers, elders and village residents. Research assistants made contact with potential study participants and made appointments for the FGDs to be conducted at a local meeting point such as office, classroom or community center. The FGDs were conducted in English for the educated class (civil servants, teachers, police officers among others) and local language among less or non educated community members. On average, the FGDs lasted one hour. Transport reimbursement of the equivalent of 3 USD was given to the participants. The transcripts conducted in local language were translated into English.

Data were collected in 11 districts that were implementing the Strengthening Civil Society for Improved HIV/AIDS (SCIPHA) and Orphans and Vulnerable children (OVC) service delivery project. The Joint Clinical Research Center implemented the SCIPHA project with funding from Civil Society Fund. The SCIPHA was a 5-year project designed to increase access and utilization of HIV/AIDS care, treatment and support services and at the same time build capacity for Civil Society Organizations to deliver quality HIV prevention, care and treatment services. The 11 districts were selected from northern and central Uganda because these two regions had the highest prevalence of HIV in the Uganda AIDS Indicator survey [21]. In central Uganda the districts were Kalangala, Mpigi, Mityana, Kiboga, and in northern Uganda they were Gulu, Lira, Amolatar, Katakwi, Moyo, Arua and Nebbi. As part of program implementation, qualitative data were collected to inform project activities.

### Data analysis

Data were analyzed using thematic approach [22]. In this method, audio recordings from the FGDs were transcribed and read carefully, back and forth. Data were analyzed manually. Texts were coded using a scheme that was prepared ahead of time. We anticipated results to fall into three broad themes of gender issues, socio-cultural factors and economic. However, emerging themes were allowed and we included attitude as a new theme. The emergence of the themes was allowed to let the data inform the analysis. The emergent codes included mobile phone, internet and social media, television, condom beliefs, migration and urbanization.

### Ethics statement

Study participants provided written informed consent and were assured the information provided was confidential. The study was submitted and approved by the JCRC Institutional Review Board and Uganda National Council of Science and Technology.

## Results

From our findings, we first, present the structural drivers of HIV and these were divided into three categories: Gender issues, socio-cultural, and economic drivers. The fourth theme, attitudes and beliefs was considered as an individual level factor. The gender issues identified were domestic violence, male involvement in health care seeking and fertility issues. The socio-cultural drivers identified were widow inheritance, funeral practices, traditional medical and circumcision practices, stigma, discrimination, and mobile phone and internet use. The economic drivers included poverty and unemployment leading to transactional sex. The individual factors include attitudes, beliefs and alcohol consumption. We also present the motivation for engagement in these structural drivers. These have been summarized and presented in a conceptual diagram in Figure 1 below:

**Figure 1:**
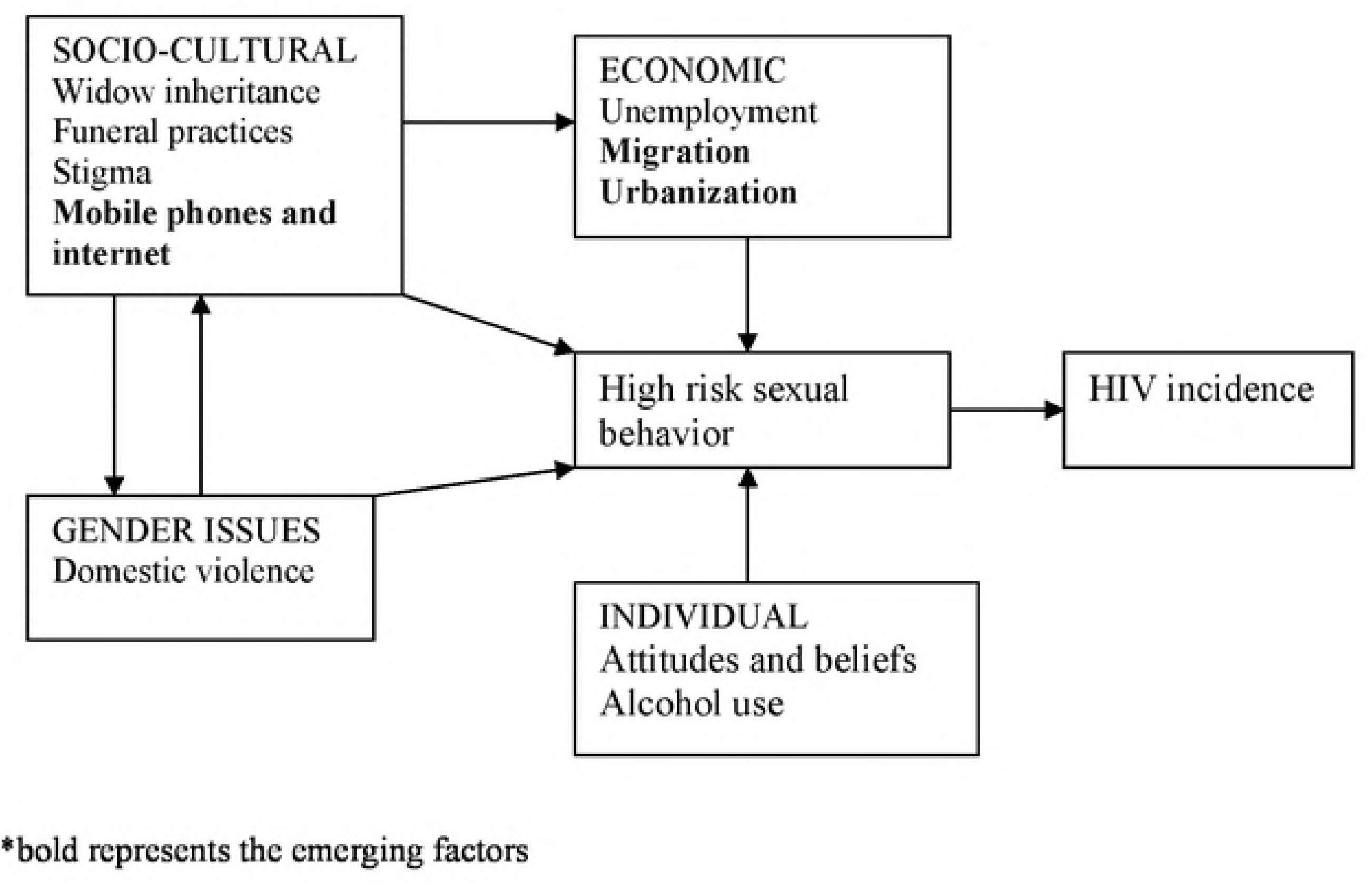
Conceptual framework to explain the traditional and emerging structural factors for HIV risk in rural Uganda.

### Gender issues

#### Domestic violence

Domestic violence was cited as one of the key factors fueling the HIV epidemic by FGD participants. Participants explained that whenever there is a misunderstanding or domestic fights within the marital union, the male partners leave the home to find new partners. Participants also mentioned that women in abusive relationships are not able to negotiate the use of condoms with their partners for fear of being abused further, are not able to seek HIV services such as HIV testing, as one participant mentioned; “*You cannot ask your husband to go for an HIV test or use a condom with you when you just had a fight. He will become even more suspicious and end up beating you more…*” FGD-22, female participant Katakwi district

Some participants attributed the problem of domestic violence to dowry and bride price stating that some women felt trapped when the relationship turned sour and were not able to leave because of the bride price that had been paid to their families.

Women commonly seek permission from their husbands before they can access health care. FGD participants mentioned that when some men discover they are HIV positive, they prevent their wives from seeking HIV testing services or even go for services for the prevention of mother to child transmission of HIV for fear of their HIV status being discovered. As one participant mentioned; “*I think they (men) fear that if the woman tests HIV positive, then she will know where the virus came from…*” FGD-33 Female participant in the island district of Kalangala.

Infertility within a couple is often blamed on the woman. The blame places pressure on her to seek for treatment from traditional sources. The study participants mentioned that traditional healers often take advantage of the women and seek to have sexual intercourse with them as part of the remedy for or treatment for infertility.

### Socio-cultural drivers

#### Widow inheritance

Widow inheritance is a common practice among some tribes in Uganda. When a woman looses her husband, the family allows the woman to select a new husband from among the brothers or relatives of the deceased or the family selects one of the relatives of the deceased to inherit the widow as their wife. When the family selects the husband to inherit the widow, the selection is done without consideration of the age differences. In our study districts, the practice was described as common in northern Uganda and west Nile region. Among the *Acholi* in northern Uganda, it is referred to as “*Lako dako*” and among the *Lugbara* in west Nile region, it referred to as “*Okoviza*” which can be literally translated as “bringing the widow back into the clan”. The inheritance typically proceeds without knowledge of the HIV status of the deceased husband or wife or even that of the new husband and therefore poses a risk of HIV transmission within the new partnership. FGD participant, Northern Uganda

> “*This is done to keep the property of the deceased within the family but also is done to maintain the family line because ….the new husband will take care of the wife and children of the deceased*”. FGD-28 Male participant, Arua, West Nile

#### Funeral practices

Following the death of a family member, several African tribes perform a ritual referred to as the last funeral rites to install a successor to the deceased. In Northern Uganda, this is referred to as “*guro lyel*”. The family of the deceased and the clan organize the function in memory of deceased member or members who have passed on during a given period of say two years. The function is organized to call the spirit of the dead back so they restore their homes “*iwongo tipu*”, to bring hope, happiness, peace and also seeking for the spirits intervention in times of sickness and material need.

The belief is that failure to hold this ceremony, the spirits of the dead will strike back and will manifest as family misfortune such as more deaths and poor harvests. During these ceremonies, local tents (*bolo*) partitioned into small rooms to accommodate visitors are constructed. It is custom that a child should be conceived during this ceremony to replace the dead. The *bolo* are designed to facilitate sexual intercourse by both youth and adults, some meeting for the first time, and in the process, exposing persons to risk of HIV acquisition.

In central Uganda, the Baganda, a majority tribe has a ritual “*Okwalula abalongo*” performed following the birth of twins, literally translated as ‘raising the twins’. The ceremony brings together relatives of the parents of the twins to celebrate the birth but also verify paternity. Participants in the FGDs indicated the ceremony may expose attendees to the risk of acquiring HIV.

> “*They engage in celebration including performance of sexually provocative dances and sing obscene songs. The ceremonies are private and often involve having sexual intercourse with strangers*”. FGD-6 Female participant, Mpigi district

In central Uganda, the FGD participants attribute the HIV epidemic to the breakdown of the cultural traditions and family ties. For instance, participants mentioned that young girls were taught good moral behaviors by their aunties or “*Ssengas”* but currently this practice has become less common. The young girls are not amenable to this advice because they do not consider it modern enough to fit the current trendy life style they lead.

In the same vein, participants mentioned the traditional family structure has broken down in several communities because of orphan hood from death of parents due to HIV leading to child headed households. Participants agreed that previously the larger family or clan would take care of the young family, but this seems to have changed because the traditional structures have been stretched. They mentioned that children now have to fend for themselves when their parents die often exposing the children to HIV:

> “*The eldest of the children takes responsibility of their siblings. They may have to drop
> out of school to find a job and this has forced some of them into prostitution ……often having sex with older men who may be HIV infected*.” FGD-11, Male participant, Kiboga district

#### Traditional medical practices

Traditional birth attendants (TBAs) provide easy access for services related to delivery and child birth in rural areas. Mothers trust the TBAs as they are members of the local community and are well known to them. However, FGD participants observed some shortcomings with the practice of some TBAs which involve use of non sterilized equipment which may result in HIV transmission:
Tattooing is traditionally practiced by placing cuts on someone’s body in order to treat an illness. FGD participants mentioned that for example when someone complains of headaches, cuts are placed on the forehead and the cuts rubbed with local herbs. The same razor blades are used to place cuts on more than one. Similarly, in the northern Uganda region, the practice of removing milk teeth of young babies aged between 4 and 6 months, referred to as “*kwanyo gi-dog*” is common. FGD participants mentioned that there is a belief that if milk teeth are not removed, they cause severe fever which may cause death. Communities have practitioners who perform these procedures and use the same unsterilized instruments including crude ones like sharp knives or bicycle spokes between patients.

Traditional medical practice also involves the use of local instruments, often non-sterilized to remove an object from the throat, “*gi dwon*”. It is done among children and the same equipment can be applied to several children without sterilization. The practice of witchcraft is widespread in the region according to reports from several in-depth interviews and FGDs. They reported that several people go to witch doctors in pursuit of wealth. For some women, their intention is to find avenues for getting a rich husband. The participants reported that witch doctors take advantage of their clients and make claims that remedies for their problems can only be delivered through sexual intercourse:
Participants believe some of these witch doctors may be responsible for a significant number of HIV infections in the community. Participants also mentioned that some witch doctors lure their HIV positive clients to stop their antiretroviral therapy with promises of better treatment options.

> “*People with HIV abandon their drugs and go for herbs and when their CD4s fall very low, they rush back to the hospital when it is too late and many of them die in the process of restarting their medications*” FGD-17, male participant, Lira district

Circumcision is traditionally practiced by some tribes in Uganda as a sign of entry in manhood. Mass circumcision of boys is conducted in traditional ceremonies and accompanying rituals are performed to initiate these boys into manhood. Participants stated that following circumcision, the boys are forced to perform some acts that may increase their risk for acquiring HIV infection.

> “*In Gishu culture, after circumcision, the boys are required to have sex with an older woman*” FGD-23, male participant, Katakwi district

#### Stigma and discrimination

The FGDs indicated the occurrence of stigma and discrimination of persons living with HIV is still widespread. They mentioned that as a result of this, that non-disclosure of HIV status is very common because of uncertainty about effects of disclosing.

> “*In some communities, people fear to share things like clothes, plates, cups or a bed and this reduces the self esteem of the HIV positive persons, so they also choose not to tell their status*”. FGD-20, Female participant, Amolatar district.

Some patients are still afraid to be seen at the HIV clinic, especially by health workers known to them and would rather seek care from a far off place, where they are not known. Study participants also mentioned that fear of disclosure was affecting decision to initiate treatment and for those who were already on treatment, their adherence was being affected by this fear.

> “*Some patients hide their medicines when visitors come around because they do not want them to know (their HIV status). But husbands are also hiding medicines from their wives and vice versa*” FGD-29, female participant, Arua district.

Domestic violence was cited as a common occurrence when disclosure happened. Women particularly expressed fear that a husband would undoubtedly engage in domestic violence if a woman disclosed she was HIV positive.

> “*There is no way I would tell my husband that I am HIV positive,that will be the end of you. The man will always think that you as a woman brought the disease in the family, even when he knows he was sick before*”. FGD-14, Female participant, Gulu district

Women were particularly concerned about the roles of their mothers-in-law. They mentioned that these in-laws are involved in their home affairs especially when there is illness. Participants were concerned that the mothers in-law ask a lot of questions about the health of their daughters in-law especially following delivery, when they even come to stay with them.

> “*There is no way a woman will swallow those medicines when the mother in law is around. You are simply inviting unnecessary question*”. FGD-26, Female participant, Moyo district.

#### Mobile phones and internet

Many youth today have access to cell phones and the internet. FGD participants agreed that these phones are being used to fuel high risk sexual behavior. The phones make it easy for the youth to share pornographic material, meet and have sex. The participants mentioned the youth now have easy access to pornographic material and can be shared easily as one participant mentioned:

> “*The students download pornographic movies onto their phones and after watching them, they want to experience what they have seen*”. FGD-10 Male participant, Kiboga district

The consensus in the FGDs was radio and television stations have also contributed in the influence of sexual behavior among young people. They mentioned that some TV and radio programs provided information that was not suitable for a young audience yet they were being aired during hours when youth are listening or watching TV. The radio station invite sexually experienced adult women (*Sengas*) to discuss these topics.

> “*The Sengas discuss sexual matters openly, discussing the most romantic styles for sex and these are picked up by the youth who want to practice and explore what has been discussed*”. FGD-8 Male participant, Mityana district

### Economic factors

#### Unemployment and economic strife

The FGD participants pointed to high rates of unemployment among the youth and participants mentioned that poverty had caused youth to become desperate and economically frustrated. Some youth will easily engage in commercial sex work to make ends meet. In places with long distance truck drivers, such as border towns, the respondents mentioned the number of commercial sex workers (CSW) had also increased.

Similarly, in eastern Uganda, FGD participants mentioned that older women recruit young girls into sex trade and act as middle men for the truckers. The participants mentioned that commercial sex work is considered illegal but seems to be on the increase as more people are getting drawn to the practice. They mentioned that single mothers have been lured into sex work in order to provide food and housing to their families and data indicated that commercial sex workers are willing to take more risk if the client offers more money.

> “*There is increased number of sex workers because of these [long distance] truck drivers and the sex workers can offer unprotected sex and charge 60,000 shillings or less at 30,000 if protected sex*”. FGD-22 Male participant, Katakwi district

Study participants mentioned that a number of younger women are choosing to have sex with much older men and vice versa in search of a more comfortable life the older partner can provide. The FGDs attributed this to the rampant poverty, families having many children and not being able to provide for them the basic needs.

> “*The young girls end up hooking older men and young boys with [older] women, who can easily provide for them*”. FGD-14, Female participant, Gulu district

#### Migration and urbanization

Migration and mobility in search of a better life and job opportunities places individuals in vulnerable positions. In our data collection, FGD participants cited examples of men who seek jobs away from home in fields such as road construction and fishing. Discussants mentioned these men earn money but are not able to spend it with their families, so they live a reckless life.

There is rapid urbanization in Uganda with proliferation of trading centers across rural areas of Uganda. Participants in the FGDs mentioned that rural areas are getting exposed to events that were commonly reserved for urban areas. For instance, discos and video halls are rapidly proliferating and provide entertainment to rural dwellers. Respondents mentioned these provide avenues for passing time but decried the potential repercussions.

> “*The karaoke discos are playing all the time and the videos show pornographic films, both day and night, children [regardless of age] spend most of their time here instead of going to school……*” FGD-14, Female participant, Gulu district, Northen Uganda

Participants in these FGDs mentioned that youth are attracted to these discos and commonly engage in sexual activities during and after these discos.

### Individual level factors

#### Attitudes and beliefs towards HIV prevention; perceived risk for HIV infection

Our data collection indicated that the way HIV prevention messages are presented to the public has changed drastically and the public now perceives HIV as a less dangerous disease compared to before as one FGD participant mentioned:

> “*In the 1980’s the HIV messages on radio were very scaring and instilled fear among people, and many were very careful not to contract HIV*.” FGD-10 Male participant, Kiboga district

The discussion also indicated that HIV disease presentation seemed to have changed and many persons with HIV/AIDS today do not look as sick as those who were infected earlier in the epidemic as one participant mentioned:

> “*….the persons with HIV looked very scaring then and nobody wished to fall in the same situation. But this has changed with ARVs. Now the disease looks like any other ailment*.” FGD-1, Male participant, Nebbi district

In some situations, individuals are aware of the risks of HIV transmission, but are not able to make the decision for safer sex because of certain attitudes. For example, in some FGDs, discussants mentioned that men often refuse to use condoms because of the attitude that sex using condoms is not pleasurable. In the discussion, participants mentioned that these men will have unprotected intercourse in high risk encounters like among commercial sex workers under this pretext. As one participant stated; “*To make the situation worse, they do not use condoms when having sex [with sex workers] saying when one buys a sweet, they do not lick it with the polythene on …*” FGD-5, Male participant, Mpigi district.

Among the fishing communities, HIV prevention is even more complex because of many other perceived competing risks. For instance, FGD participants mentioned that at the end of every fish harvest, the fishermen celebrate to mark the success. They buy alcohol and engage in sexual intercourse with random partners. They consider the lake to be much more dangerous than HIV. As one member pointed out; “*They fear water more than HIV, [which] they call a small organism that gives you a grace period to eat what you have worked for and even prepare your will, but with water, you are gone in a blink of an eye*” FGD-33, Male participant, Kalangala district

The fishing villages are dominated mostly by males, with women as a minority. Study participants mentioned that there is competition for the fewer women as sexual partners that it is very common for a woman to have multiple sexual partners, creating a concurrent sexual partnership.

The FGDs revealed that among some tribes, there is resistance to the use of condoms with the belief that certain sexual styles are not compatible with use of condoms. The discussions also showed that in some communities, there is a belief that having sexual intercourse with a person with disability (PWD) brings blessings. This places PWD in a vulnerable position and increases their risk for HIV.

#### Drug use and alcoholism

Most of the FGD participants agreed that alcohol consumption was very high and is one of the potential drivers for HIV infection among youth, both directly and indirectly. Alcohol is freely available and there are no restrictions on access based on age category or opening and closing hours for bars. In the schools, respondents mentioned that both teachers and students drink alcohol and end up in bars where they are exposed to high risk sex, and as a respondent in northern Uganda mentioned, this may be aggravated because some schools are very close to bars and disco halls.

> “*But it is worse when the teacher comes to teach and he is drunk, as pupils are caught up in laughter because of the teacher’s behavior. Students simply run out of class and go to disco halls where they hook up for engagement in sexual activities*” FGD-16, Male participant, Lira district.

Among the out of school youth, alcohol consumption is uninhibited. Participants mentioned the rampant use of other substances such as marijuana and that it is common for motorcycle taxi (*boda boda*) riders, who ride the motorcycles under the influence of these substances lure or force their customers into having sex with them.

## Discussion

Our study has indicated that several structural factors and drivers of HIV exist in rural Uganda, and are of a varied nature. These factors range from gender issues, social cultural, traditional medicinal practices to economic factors. Our data indicates these factors have persisted despite decades of HIV prevention. Some of these factors have previously only been anecdotally reported. Our paper has explored these traditional structural drivers but also sheds light on new ones that have resulted from urbanization and modernization such as exposure to mobile phones and the internet. These drivers are connected and intertwined in a complex web as illustrated in our conceptual framework. However, they provide a basis for integrated structural interventions, taking into consideration traditional HIV prevention, care, treatment issues. Our paper also makes recommendations useful for the design of these new interventions that may form the core of the next generation of HIV prevention programming in order to achieve an HIV/AIDS free generation.

Our data shows significant homogeneity across the country in terms of drivers. Regardless of the region, the factors that influence HIV appear to be the same. Previously, other studies have documented structural factors and drivers of HIV infection and results indicate some homogeneity and heterogeneity within and between communities, tribes and countries among these drivers [8, 12]. Our study also shows the same in comparison to these studies. For instance, while as transactional sex was common in our study and several others, there are some peculiar practices such as “*kusomboka*” or death cleansing which includes unprotected sexual intercourse with a corpse which have been reported only in Tanzania[23]. This finding implies that design of interventions will require an understanding of the local drivers.

Our study also reveals the factors have a highly gendered stratification placing women at a disproportionately higher risk for HIV infection compared to men. For instance, in our data collection, women were reported as more likely to face higher risk for HIV infection in situations of domestic violence, widow inheritance, last funeral rites and traditional medical practices to treat female infertility. In India, illiteracy of the woman and early marriage have been shown to be major drivers of HIV infection in a multi state study in the south of the country [24].

In our study, participants indicated that stigmatization of HIV patients was still common. The results on stigma have been conflicting. Some participants mentioned the availability of treatment had ‘normalized’ the disease to appear as any other. The data suggesting a reduction in stigma agree with those from a recent study in Uganda which showed there has been a reduction in self stigmatization due to availability of ART[25], however in this study, the structural drivers of stigmatization such as gender inequalities were still common and were a barrier to elimination of stigma. Another recent study in rural Uganda [26] has shown stigma may persist despite availability of antiretroviral therapy, signifying stigma remains a structural barrier to deal with.

Commercial sex work was mentioned as a common practice and was largely driven by economics. A recent study in Malawi [27] showed the structural drivers of commercial sex work among young adults were deprivation of food, housing and a desire for trendy items like clothing and cellular phones. The participants in our study mentioned these same factors drive young single mothers into commercial sex work. Although not mentioned in our FGDs, other studies have found young people are also at risk for coercive sex if they are from materially deprived households [28]. Several other studies across Africa [29–31] have also showed a relationship between material deprivation and high risk sexual behavior. Interventions for HIV prevention among vulnerable groups should also include programs for economic empowerment and job creation to reduce material deprivation and unemployment. As other authors have suggested, an integrated approach to HIV prevention incorporating gender and development programs is a necessity [32]

Mobility and migration have been identified as structural drivers for HIV transmission in Tanzania [23] among a fishing population and in Uganda [33], among the youth. In our study, significant mobility was noted among the fisher folk. Similar to the Tanzania study, our findings among a fishing population indicate the same finding. Mobility is difficult to ta

Interventions with the participation of local entities like local governments, traditional and cultural leaders should be designed to target these structural drivers. Studies need to be done to rigorously test feasible and promising interventions such as one to keep adolescent girls in school in India [34], Safe Homes and Respect for Everyone (SHARE) project in Uganda, a intervention against domestic violence which has shown a reduction in physical and sexual intimate partner violence and a corresponding reduction in HIV incidence [35, 36]. Another such project, SASA!, also in Uganda showed community level interventions can result in reduction of intimate partner violence, high risk sexual behaviors and strengthening community based structures to enhance these changes [37–39].

Evidence to suggest that structural interventions may be the solution has also been observed in rural Uganda, where reduction in HIV incidence among adolescent girls was attributed to increase enrollment in school [40]. Such interventions will need to be scaled up to realize sustained decreases in HIV incidence.

## Conclusion

In conclusion, our study has outlined several structural drivers that have for long been anecdotal, but also verifies other well known drivers. Our data indicates these drivers have persisted in the communities despite decades of HIV prevention. Many of these drivers are very complex and intimately intertwined with cultural heritage and local traditions and need to be tackled cautiously with knowledge of the modern and promising interventions.

## List of abbreviations

FGD: Focus group discussions
JCRC: Joint Clinical Research Center
OVC: Orphans and Vulnerable children
SCIPHA: Strengthening Civil Society for Improved HIV/AIDS
TBA: Traditional birth attendant

## Declaration

### Ethics approval and consent to participate

Study participants provided written informed consent and were assured the information provided was confidential. The study was submitted and approved by the JCRC Institutional Review Board and Uganda National Council of Science and Technology.

### Availability of data and materials

Detailed data and transcripts are available on request, after appropriate permissions have been obtained.

### Authors’ Contributions

FT, CMK, GA, PNM conceived the idea. FT, GA, DA, FB collected the data, transcribed the discussions, coded and FT, GA, DA, FB, CMK, PNM analyzed the data, agreed on structure of draft of manuscript. FB, GA, FT drafted the first draft. All authors revised the manuscript and agreed on the final version for publication.

## Acknowledgements

The authors would like to thank the SCIPHA field team and these include the district site and regional coordinators, data managers and the drivers for the assistance in organization of setting appointments, focus group discussions and data collection. We thank the District Health officers in SCIPHA districts for the assistance in making data collection smooth. The authors thank Civil Society Fund for supporting this study. The funder has no role in the design and interpretation of these study results.

